# CASCADE-Cas3 Enables Highly Efficient Genome Engineering in *Streptomyces* Species

**DOI:** 10.1101/2023.05.09.539971

**Authors:** Christopher M. Whitford, Peter Gockel, David Faurdal, Tetiana Gren, Renata Sigrist, Tilmann Weber

## Abstract

Type I CRISPR systems are widespread in bacteria and archaea. The main differences compared to more widely applied type II systems are multi-effector CASCADE needed for crRNA processing and target recognition, as well as the processive nature of the hallmark nuclease Cas3. Given the widespread nature of type I systems, the processive nature of Cas3, as well as the recombinogenic overhangs created by Cas3, we hypothesized that Cas3 would be uniquely positioned to enable efficient genome engineering in streptomycetes. Here, we report a new type I based CRISPR genome engineering tool for streptomycetes. The plasmid system, called pCRISPR-Cas3, utilizes a compact type I-C CRISPR system and enables highly efficient genome engineering. pCRISPR-Cas3, outperforms pCRISPR-Cas9 and facilitates targeted and random sized deletions, as well as substitutions of large genomic regions such as biosynthetic gene clusters. Without additional modifications, pCRISPR-Cas3 enabled genome engineering in several *Streptomyces* species at high efficiencies.

## Introduction

### Large diversity of unexplored BGCs, large untapped potential

Streptomycetes and other filamentous actinomycetes represent some of the most gifted producers of complex specialized metabolites, with a wide diversity of applications in medicine, industry, and agriculture. Most importantly, many of the commonly used antibiotics are produced by members of the genus *Streptomyces* ^1^.

Many of these specialized metabolites play important ecological roles for the producer strains, to fend of competition for nutrients, communication, as well as for complex symbiotic relationships ^2^. Due to the complex role specialized metabolites play in the natural environment, the expression of the corresponding biosynthetic gene clusters (BGC) is tightly regulated, and natural elicitors often remain unknown. Therefore, a majority of BGCs is not readily expressed under standard laboratory conditions, hindering the identification of the encoded specialized metabolites.

While classical chemical screening resulted in the identification and utilization of high numbers of readily expressed BGCs, the advent of cheap whole genome sequencing revealed a large untapped potential of silent BGCs encoding the biosynthetic information for yet unknown specialized metabolites ^3, 4^.

### Expression of BGCs in engineered heterologous host’s common practice

To overcome these limitations and to advance the exploration of the incredible biosynthetic potential of streptomycetes, several methods were developed to achieve expression of silent BGCs. Expression of silent BGCs in a heterologous host represents one of the commonly used approaches ^5, 6^. Since many actinomycetes are not (yet) genetically tractable, expression of BGCs cloned from isolated genomic DNA allows the study of BGCs independently of their source. Furthermore, using established expressions hosts streamlines experimental work, as no new methods need to be developed and each experiment has the potential to contribute to a growing knowledgebase. Commonly used hosts for expression of heterologous BGCs include various derivatives of *S. coelicolor*, *S. albidoflavus*, *S. lividans*, *S. avermitilis,* and *S. venezuelae* ^7–15^.

### Genome Reduction desired for reduced metabolic background activity

To increase the success rate of heterologous expression of BGCs, hosts are usually engineered with focus on three aspects. First, to aid the identification of compounds in metabolomics analysis, a simplified metabolic background is desired. This can be achieved through genome reduction with special focus on native, readily expressed BGCs, as well as those usually silent under laboratory conditions. Second, to increase the chances of successful heterologous expression, the number of integration sites can be increased, to facilitate multicopy expression of the target BGC. Lastly, increasing the supply of precursors is also of interest to boost production of a matching BGC. Examples of hosts constructed using these principles are *S. albidoflavus* BE4, *S. lividans* ΔYA11, or *S. avermitilis* SUKA22 ^7, 8, 14, 16^.

### Common methods to achieve deletions PCR targeting/homologous recombination suicide vectors/ Cas9, Cas12a

A wide number of tools exist that can be used to achieve above mentioned host engineering ^17^. Classically, genomic deletions were achieved through PCR targeting, which is based in homologous recombination and double crossovers ^18^. While this method works in many *Streptomyces* strains, it requires to have the editing site cloned in a cosmid, fosmid or BAC and thus still is very labor and time intensive. Newer methods usually make use of CRISPR effectors, such as Cas9 or Cas12a. Several CRISPR-Cas9 based tools were developed for application in streptomycetes and were shown to facilitate small to medium sized deletions with good efficiencies. Common vectors include pCRISPomyces, pCRISPR-Cas9, or pKCcas9dO^19–21^. Frequently observed toxicity of Cas9, especially when strongly expressed, has since led to the development of optimized plasmids allowing tighter control of the expression of Cas9 ^22^. pKCCpf1 was developed to enable Cas12a (Cpf1) based deletions of one or two genomic loci, based on crRNA processing by Cas12a of its own crRNA array ^23^. Cas12a further recognizes a T-rich PAM sequence, lowering the potential of off-target effects in the GC rich streptomycete genome. Recently, base editing was developed for single and multiplexed inactivation of genes of interest through introduction of premature stop codons ^24^. While this method has many benefits, the targeted genes remain present in the genome, and off-targets are more tolerated due to the absence of DSB introduction.

### CASCADE Cas3, natural function, properties

Type I CRISPR systems are among the most widespread CRISPR systems. The signature effector of type I systems is the nuclease Cas3, which requires the assembly of a ribonucleoprotein complex called CASCADE, usually formed by Cas8, Cas7, Cas5, and the crRNA, to bind and cut DNA ^25^. Hence, CASCADE Cas3 is a multi-effector CRISPR system which differs significantly from single effector CRISPR systems like Cas9. The most widely studied type I systems are type I-E, of which the *Escherichia coli* system has been studied in great detail ^26, 27^. Type I and II CRISPR systems not only differ in their overall architecture, but also in the mode of action of the signature nuclease. While Cas9 acts only as an endonuclease by introducing double strand breaks at the target site, Cas3 has a combined ATP-dependent nuclease-translocase activity and starts degrading DNA processively from 3’ to 5’ after cutting^28^. This results in the formation of long 5’ overhangs. CASCADE Cas3 has previously been applied for genome engineering, but the large size of the complex has hindered widespread application using plasmid based systems ^29, 30^. Recently, Csörgő et al. described a compact type I-C system and its application for plasmid-based genome engineering in *E. coli*, *Pseudomonas syringae* and *Klebsiella pneumoniae* ^31^. The described system requires only 4 genes (*cas3, cas5, cas8,* and *cas7*) to introduce efficient genomic deletions of up to half a megabase. The system uses a T-rich 5’-TTC-3’ PAM sequence, making it an attractive system to use in high GC organisms. Furthermore, it was shown that homology directed repair in combination with CASCADE Cas3 introduces genomic modifications with higher efficiencies than Cas9, likely due to the recombinogenic nature of the ssDNAse activity of Cas3.

### Our tool and applications, need for efficient deletion of large BGCs

Here, we present a new CASCADE Cas3 based tool for streptomycetes facilitating highly efficient genomic deletions and integrations based on a previously published compact type I-C CRISPR system. We adapted the system for expression in streptomycetes and integrated it into our established CRISPR plasmid platform. We demonstrate highly efficient deletions of small, mid, and large sized deletions, and show that pCRISPR-Cas3 can facilitate deletions in several commonly used *Streptomyces* hosts. We show how pCRISPR-Cas3 can facilitate simultaneous deletions and integrations with superior efficiencies, allowing streamlined genome engineering in even recalcitrant *Streptomyces* strains. Finally, we demonstrate the application of genome engineering with pCRISPR-Cas3 by construction of a *S. coelicolor* expression host.

## Results

### Distribution of Type I CRISPR Systems in Streptomycetes

Type I CRISPR systems are widespread in bacteria and archaea. However, only few systems from *Streptomyes* have been characterized and described in detail so far ^32^. To investigate the distribution of type I CRISPR systems vs. type II CRISPR systems in streptomycetes, we performed BLAST searches against 2401 high quality publicly available *Streptomyces* genomes. The amino acid sequence of Cas9 from *Streptococcus pyogenes* and of Cas3 of the type I-E CRISPR system of *S. albidoflavus* (previously *albus*) J1074 were used as queries, as these are the hallmark nucleases from the two CRISPR types. Based on our search, type I CRISPR systems appear to be much more widespread in *Streptomyces* compared to type II CRISPR systems. Only two hits were obtained for *Spy*Cas9, while *Salb*Cas3 returned almost 1400 hits (Fig. 1). Of these, over 100 had a sequence similarity >50 % (Supplementary Information Table 1). Cas3 acts as a nuclease-translocase with a N-terminal HD phosphohydrolase and C-terminal helicase domain ^33^. To obtain a more granular view of the distribution of type I systems in streptomycetes, and to reduce the number of false positive hits due to similarities to helicases, we performed another search with cblaster, which allows the search for clustered homologous sequences ^34^. Using cblaster and the CASCADE from *S. albidoflavus* J1074 as input, we identified 472 strains carrying the entire CASCADE, comprising of *cas3, casA, casB, cas7, cas5, and cas6* (Supplementary Information Table 2). Based on the high abundance of type I CRISPR systems in streptomycetes, we hypothesized that they might have a higher tolerance to genome engineering using type I CRISPR systems, as opposed to type II systems.

**Figure 1:**
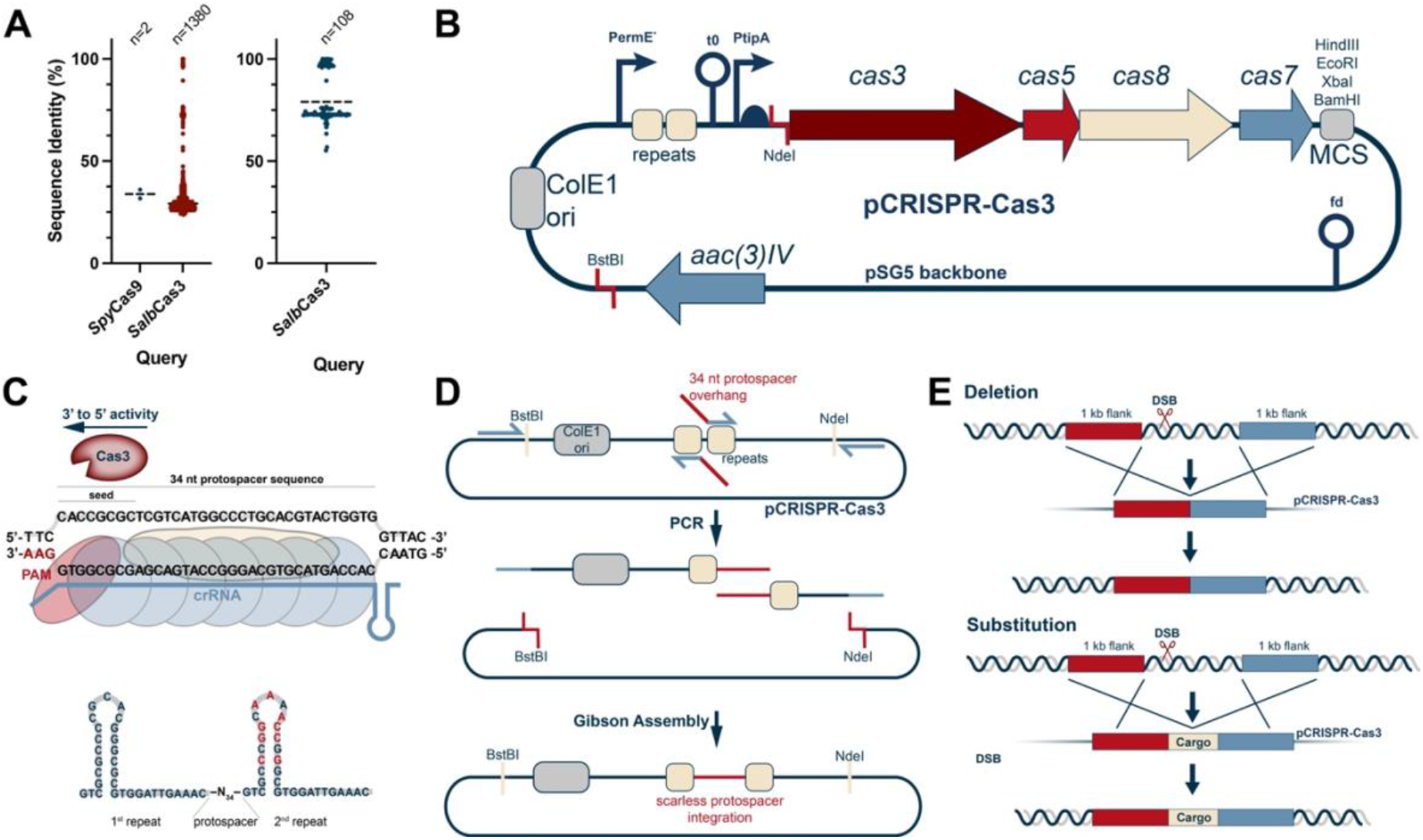
pCRISPR-Cas3 enables streamlined genome engineering of streptomycetes. **(A)** Distribution of type I and type II CRISPR systems in streptomycetes using *Spycas*9 and *cas*3 from *S. albidoflavus* as references. The BLAST search was run on all high quality publicly available *Streptomyces* genomes (n=2401). Type I CRISPR systems appear to be much wider distributed than type II CRISPR systems in streptomycetes. **(B)** Plasmid map of pCRISPR-Cas3. The plasmid is based on the pSG5 replicon and carries the codon optimized type I-C minimal CASCADE under control of the inducible *tip*A promoter. The crRNA is cloned between to repeats in the cRNA cassette, which is controlled by the constitutive *erm*E* promoter. Repair templates are cloned into a MCS in the backbone of the plasmid. **(C)** The second repeat has a modified sequence to prevent recombination between the two repeats. The CASCADE complex comprised of Cas5, Cas8, and Cas7 units binds the target sequence and recruits Cas3. Cas3 has a 3’-5’ helicase nuclease activity, resulting in directionally biased deletions. **(D)** cRNAs are cloned between two repeats, posing some challenges due to sequence homologies. Since type IIS restriction enzymes cannot be used in high GC *Streptomyces* plasmids, a PCR and Gibson Assembly based cloning approach was established, allowing cloning of cRNAs with high efficiencies. **(E)** pCRISPR-Cas3 can be used for targeted deletions of large genomic regions, or for substitutions of such with a specified cargo. It can also be used for random sized deletion experiments.

### Construction of pCRISPR-Cas3, a plasmid based compact CASCADE-Cas3 system

The previously characterized and utilized type I-C CRISPR system from *Pseudomonas aeruginosa* consists of only four genes, *cas3, cas5, cas8,* and *cas7,* totaling 5.6 kb. This allows plasmid-based expression of the system and use across various species. All four genes are arranged in an operon and can hence be expressed from a single promoter. Given the differences in both codon usage and GC content, the previously published operon was codon optimized and synthesized. For codon optimization, the *S. coelicolor* codon usage table was used, as such optimized constructs have previously been successfully expressed in a wide variety of different streptomycetes ^35^. The upstream region of each *cas* gene was designed with a canonical RBS sequence (GGAGG or GGAGC). The 5.6 kb fragment was subsequently synthesized and delivered as a cloned plasmid. For expression in streptomycetes, our established CRISPR platform based on the pSG5 replicon was used. The Cas9 cassette from pCRISPR-Cas9 was removed by digestion with NdeI and HindIII and replaced by the synthesized CASCADE-Cas3 operon using Gibson Assembly. The Cas9 gRNA expression cassette was subsequently replaced by a dsDNA fragment containing the Cas3 repeats following digestion of the plasmid with NcoI and SnaBI and subsequent Gibson Assembly. The second repeat was modified as described by Csörgő et al. to prevent recombination events between the two repeats (Fig 1c). The resulting plasmid was named pCRISPR-Cas3 in accordance with our previous plasmid nomenclature.

Given the repetitive nature of the Cas3 crRNA hairpins, cloning spacer sequences in between these proved to be very challenging. Attempts to clone protospacer sequences using ssDNA oligo bridging by digestion of pCRISPR-Cas3 with NcoI (cutting between the two repeats) failed repeatedly. Given that common type IIS restriction enzymes such as BsaI and. BstBI, commonly used for such cloning scenarios, cannot be used in high GC plasmids due to their omnipresent recognition sequences further complicated this cloning challenge. To achieve efficient cloning of user defined protospacer sequences between the CRISPR repeats, we hence had to design an alternative cloning strategy. Using two fixed primers binding up and downstream from the protospacer integration site, two PCRs are set up using two primers binding one of the repeats each and carrying the desired 34 nt protospacer sequence in the overhangs. Digestion of pCRISPR-Cas3 with BstBI and NdeI removes the origin of replication, ensuring that only correctly assembled plasmids can replicate. Using this cloning method, we frequently achieved over 90 % cloning efficiencies (Supplementary Information Figure 1).

### pCRISPR-Cas3 introduces highly efficient deletions with single nucleotide precision in S. coelicolor

To demonstrate the application of pCRISPR-Cas3 for genome engineering in streptomycetes, we attempted to delete the well characterized 22 kb actinorhodin BGC in *S. coelicolor*. Deletion of the actinorhodin BGC leads to a clear red phenotype in *S. coelicolor*, resulting from expression of the undecylprodigiosin BGC and enabling phenotypical screening for deletion mutants. Three different protospacers were selected, in the middle and towards each edge of the deletion region, to investigate potential differences in editing outcomes based on protospacer positions. Protospacers for pCRISPR-Cas3 were manually designed based on PAM+seed sequence queries within the deletion region, followed by genome wide queries using both the seed and whole length sequences of putative protospacers to identify the most specific candidates. In *S. coelicolor,* 158,341 5’-TTC-3’ PAM sequences were found, averaging one PAM for every 54.7 bp. In contrast, 1,574,641 5’-NGG-3’ PAM sequences were found for Cas9, averaging one for every 5.5 bp, and highlighting the presence of up to an order of magnitude more potential off-target sites. Two flanking regions of 1 kb were selected as repair templates and cloned into pCRISPR-Cas3 using restriction cloning as described in the methods section. To compare editing outcomes to those achieved with Cas9, the same repair templates were cloned into pCRISPR-Cas9, and three different sgRNAs mirroring the positions of the Cas3 protospacers were selected. In parallel, all experiments were performed without repair templates for both pCRISPR-Cas3 and pCRISPR-Cas9.

Stark differences were observed between the different editing configurations. pCRISPR-Cas9 without repair templates failed to produce the distinct red phenotype, and with the “right” protospacer, only a handful of viable exconjugants were obtained on selective medium. Given that the pCRISPR-Cas9 plasmid without ScaLigD was used, higher toxicity was expected. As shown previously, coexpression of ScaLigD can greatly enhance NHEJ and reduce the toxicity in *S. coelicolor* ^20^. pCRISPR-Cas3 without repair templates resulted in greatly varying outcomes, depending on the used protospacer. Both protospacers close to the edges of the deletion region appeared to be toxic. However, both surviving clones of the right protospacer had a weak red phenotype. pCRISPR-Cas3 without repair templates and the protospacer in the middle of the deletion region did not result in observable loss in viability, but also did not produce a clear red phenotype. Clear deletion phenotypes were obtained with pCRISPR-Cas9 with repair templates and the spacer located in the middle of the deletion region. However, with the other two protospacers, either no clear deletion phenotypes were obtained, or only a small number of viable exconjugants could be obtained, which did not have the desired phenotype. The observed toxicities of pCRISPR-Cas9 might be predominantly protospacer specific. However, recent studies have shown that binding to off-target sites might be much more extensive, stabilized by as little as a few nucleotides of homology ^36^. Using pCRISPR-Cas3 together with repair templates, clear deletion phenotypes were obtained for all spacers. With protospacers located towards the edges of the deletion region, clear deletion phenotypes were obtained for all picked exconjugants. Only 5 out of 12 exconjugants showed the expected phenotype when the protospacer in the middle of the deletion region was used, with the remaining 7 having a greyish phenotype,. Interestingly, these results appeared to be mirrored to pCRISPR-Cas9, which performed best with the protospacer located in the middle of the deletion region.

Phenotypical screening revealed an apparent superiority of pCRISPR-Cas3 compared to pCRISPR-Cas9, as all three protospacer configurations resulted in the expected deletion phenotype with high efficiencies. To verify these observations and to obtain more precise numbers for efficiencies of pCRISPR-Cas9 and pCRISPR-Cas3, the experiments with protospacers and repair templates were repeated. This time, conjugations were performed at larger scales to obtain more viable clones for pCRISPR-Cas9 with the protospacer located in the right flank of the deletion region. The deletion efficiencies were analyzed by performing colony PCRs on the targeted region with a primer binding inside of the repair template, and one binding outside of the repair template in the neighboring sequence (Supplementary Information Figure 2). For pCRISPR-Cas9, average deletion efficiencies of 60 % were obtained. These varied greatly depending on which protospacer was used. For pCRISPR-Cas3, an average efficiency of 97 % was obtained. The obtained efficiencies for pCRISPR-Cas3 were consistently high for all protospacers, suggesting a lower dependency on the protospacer sequence and location to achieve efficient deletions.

**Figure 2:**
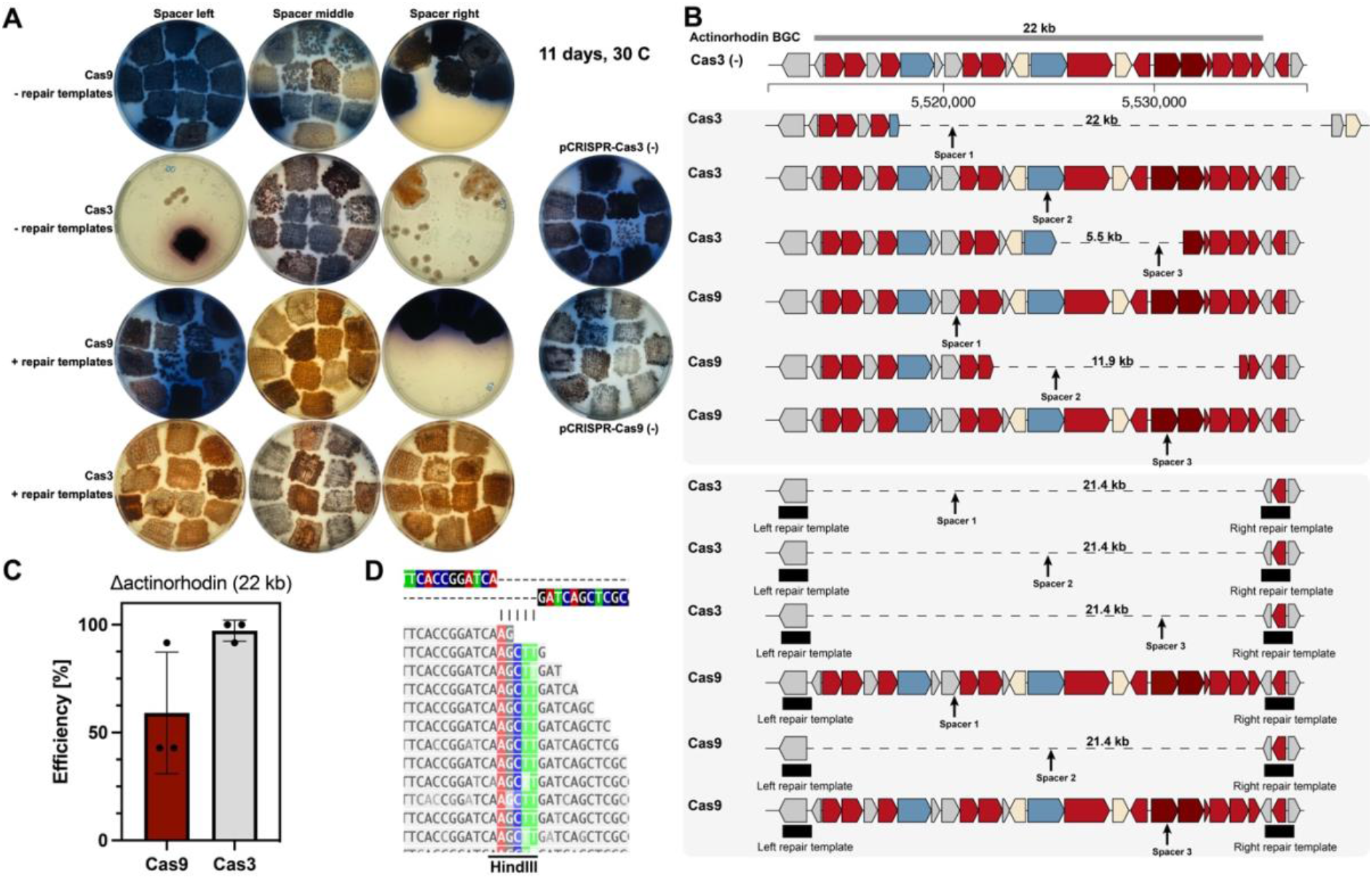
pCRISPR-Cas3 introduces genomic deletions with higher efficiencies than pCRISPR-Cas9. **(A)** Plate pictures of *S. coelicolor* mutants harboring pCRISPR-Cas9 or pCRISPR-Cas3 with protospacers targeting the actinorhodin BGC in three different locations, and with or without repair templates. pCRISPR-Cas3 displayed higher toxicity without repair templates, but resulted in more exconjugants overall and with the desired RED phenotype once repair templates were provided. **(B)** Representation of sequencing results of selected colonies, both for pCRISPR-Cas3 and pCRISPR-Cas9 with and without repair templates. Both pCRISPR-Cas3 and pCRISPR-Cas9 introduced random sized deletions without repair templates. With repair templates, precise deletions were observed for both pCRISPR-Cas3 and pCRISPR-Cas9. (**C)** Efficiencies for actinorhodin deletions with pCRISPR-Cas9 and pCRISPR-Cas3. For pCRISPR-Cas3, the efficiencies were consistently high (>90 %), while with pCRISPR-Cas3 the observed efficiencies were highly dependent on the used protospacer. **(D)** Read alignments for the junction site of the two homologous flanks. A HindIII site was integrated, demonstrating that the double strand break was repaired using the repair templates cloned into pCRISPR-Cas3 using HindIII. Shown in (**C**) are the means ± standard deviations of three deletion experiments targeting the actinorhodin region with three different protospacers.

To verify the PCR results, Illumina based whole genome sequencing was performed for one colony of each configuration (Fig. 2B). Random sized deletions were observed for both pCRISPR-Cas9 and pCRISPR-Cas3 when no repair template was provided. For pCRISPR-Cas3, the observed random sized deletion were 5.5 kb and 22 kb in size, and for pCRISPR-Cas9 11.9 kb. No random sized deletions were observed for pCRISPR-Cas3 with the mid protospacer, and none for pCRISPR-Cas9 when the protospacer at the edges of the deletion region were used. Sequencing of repair template containing configurations showed that pCRISPR-Cas3 successfully introduced the designed mutations with single nucleotide prevision with all tested configurations. For pCRISPR-Cas9, only the protospacer located in the middle of the deletion region resulted in successful deletion of the designed region for the screened colonies.

As pCRISPR-Cas3 was able to introduce the designed deletions with robust efficiencies and single nucleotide precision, we next investigated whether pCRISPR-Cas3 could be used to install targeted deletions in other important *Streptomyces* species.

### Genomic deletions were achieved using pCRISPR-Cas3 in *S. venezuelae, S. albidoflavus, S. sp.* NBC01270

Given the successful demonstration in *S. coelicolor*, we next attempted to install designed deletions in other strains of interest. Therefore, we selected *S. venezuelae* (ATCC 10712), an emerging production host and model species, *S. albidoflavus* J1074, a well-established host for expression of heterologous BGCs, and *S. sp.* NBC1270, an isolate from our own strain collection closely related to *S. albidoflavus* J1074.

In *S. venezuelae* ATCC 10712, region 22, as predicted by antiSMASH 6.0.1, was selected as a target for demonstration, given its size of 122 kb, and the high density of putative BGCs. Repair templates of 1 kb on each side of the target region were designed. Two protospacers were designed, one in the middle of the deletion region, and one approximately 5 kb from the left edge. Both combinations of repair template and protospacer installed the desired deletion with high efficiencies, with 9 out of 12, and 10 out of 12 exconjugants carrying the designed mutation, respectively (Fig. 3). The PCR based screening results were further verified by whole genome sequencing, showing a clear drop in coverage of region 22.

**Figure 3:**
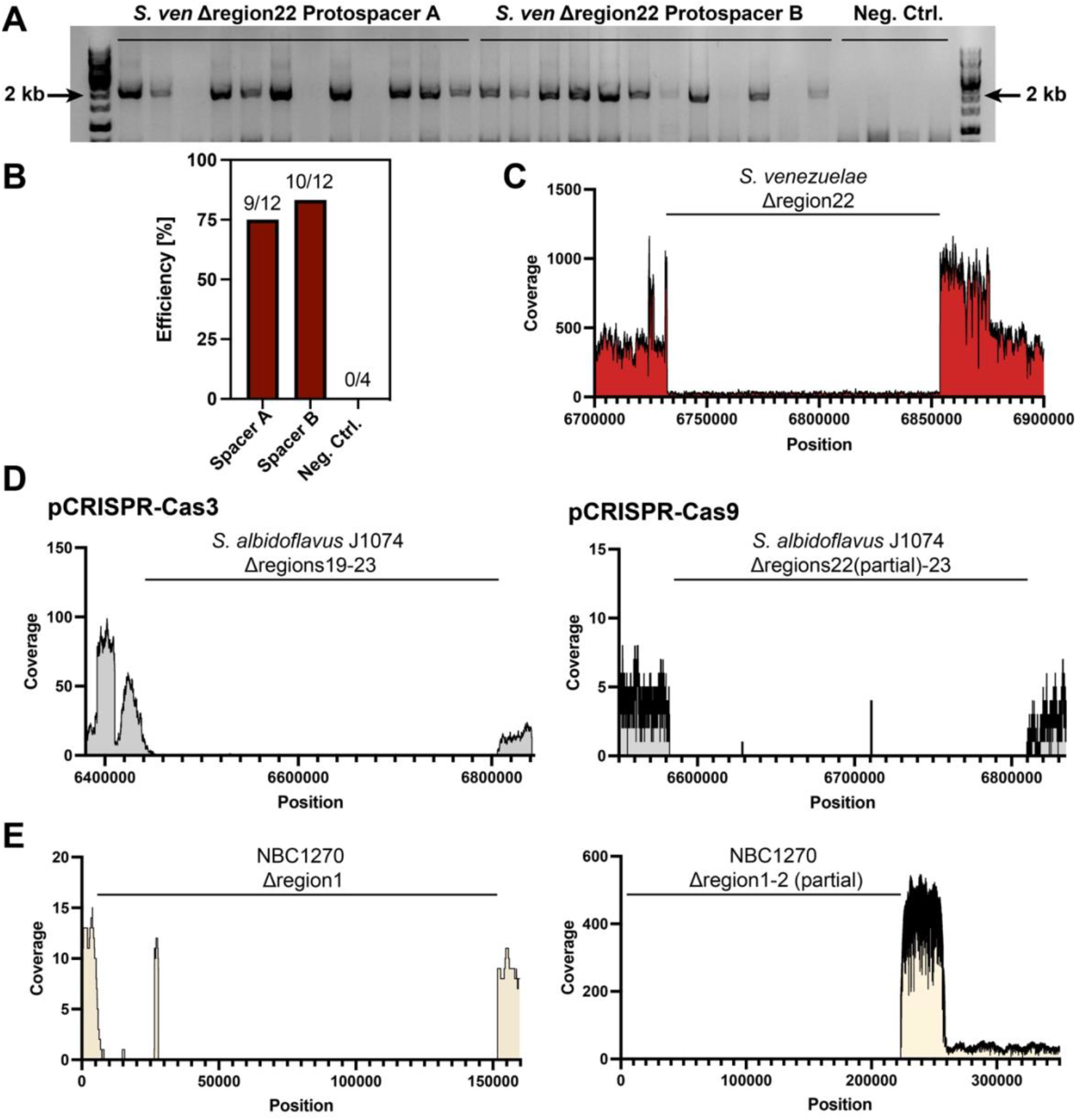
Application of pCRISPR-Cas3 in other *S. venezuelae, S. albidoflavus* J1074, and *S. sp.* NBC1270. **(A)** Gel picture for screening of successful deletions of the 122 kb region 22 in *S. venezuelae*. Bands were expected slightly above the 2 kb marker, as the primers were binding just outside of the repair templates. **(B)** pCRISPR-Cas3 was able to introduce the desired deletions with high efficiencies for both protospacers. **(C)** Illumina based sequencing of a deletion mutants showing a clear and precise deletion of the targeted region. **(D)** In *S. albidoflavus* J1074 random sized deletions were detected following targeting of regions 21-23 with pCRISPR-Cas3. However, similar random sized deletions were obtained when targeting the same region with the same repair templates using pCRISPR-Cas9. **(E)** In *S. sp.* NBC1270, a *Streptomyces* strain with high similarity to *S. albidoflavus* J1074, random sized deletions were obtained with pCRISPR-Cas3 while preserving the terminally inverted repeats. With pCRISPR-Cas9, targeting of the same region resulted in random sized deletions and complete loss of the chromosomal end. A 33 kb sequence likely got replicated up to 12 times to prevent continuous degradation of the chromosomal end.

Given the positive results in *S. venezuelae*, we next attempted to perform deletions in *S. albidoflavus* J1074 and the closely related strain *S. sp.* NBC1270. The strains are closely related and share the majority of BGCs as predicted by antiSMASH 6.0.1. At the far end of the chromosomal arm, both strains carry the same dense accumulation of 3 BGCs for production of antimycin, candicidin, and flaviolin. Given the total size of 318 kb, this BGC dense region was also interesting in order to investigate the possibility to delete hundreds of kb in a single step. Following several failed attempts to obtain the bands indicating successful introduction of the designed chromosomal deletion, some of the colonies for which no PCR bands were obtained were sequenced using Oxford Nanopore sequencing. In both *S. albidoflavus* J1074 and *S. sp.* NBC1270, targeting of the far chromosomal end resulted in unspecific deletions approximately 380 kb and 140 kb in size, respectively. In *S. albidoflavus* J1074, this resulted in deletion of antiSMASH regions 19-23. Targeting of the same region with pCRISPR-Cas9 also resulted in extended deletions, suggesting that this was a general problem resulting from introducing double-strand breaks at the ends of the chromosomal arms ^37^. Interestingly, targeting the same region in NCB1270 using pCRISPR-Cas9 resulted in loss of the entire chromosomal end, as no reads were mapped against the terminally inverted repeats. A big spike in coverage was observed just at the end of the deletion region, suggesting that this 33 kb sequence stretch was replicated around 12 times to prevent continuous degradation of the chromosomal end. De novo assembly of the genome and subsequent visualization of the assembly graph further confirm this hypothesis (Supplementary Information Fig. 3). Both strains did not display any growth defects, and all desired BGCs were deleted in *S. albidoflavus* J1074, despite the unspecific nature of the deletion. These results demonstrate that pCRISPR-Cas3 can be used to introduce random sized deletions, especially when coupled with subsequent screening for desired geno- and phenotypes. Our results highlight that pCRISPR-Cas3 can enable efficient genome engineering even in non-model *Streptomyces* species.

### Simultaneous deletions and integrations can be achieved through modification of the repair templates

Most sophisticated engineered *Streptomyces* hosts have not only a reduced metabolic background, achieved through deletion of readily expressed BGCs, but also added integration sites for either multicopy integrations or site directed targeted integrations using different integrases ^7, 38, 39^. Consequently, we wanted to demonstrate how pCRISPR-Cas3 can be used to delete genomic regions such as BGCs, and to simultaneously integrate additional integration sites instead. The *Streptomyces* bacteriophage PhiC31 integrase is as well-established system for integration of heterologous sequences in various *Streptomyces* species ^40, 41^. The consensus PhiC31 attB site from *S. coelicolor*^42^ was chosen as cargo and cloned it in between the two flanks. As a proof of concept, we modified plasmid p129, previously used to delete the actinorhodin BGC in *S. coelicolor,* to carry the attB site between the two homologous repair template arms. The obtained sequencing reads mapped perfectly against the *in silico* modified reference sequence (Fig. 4).

**Figure 4:**
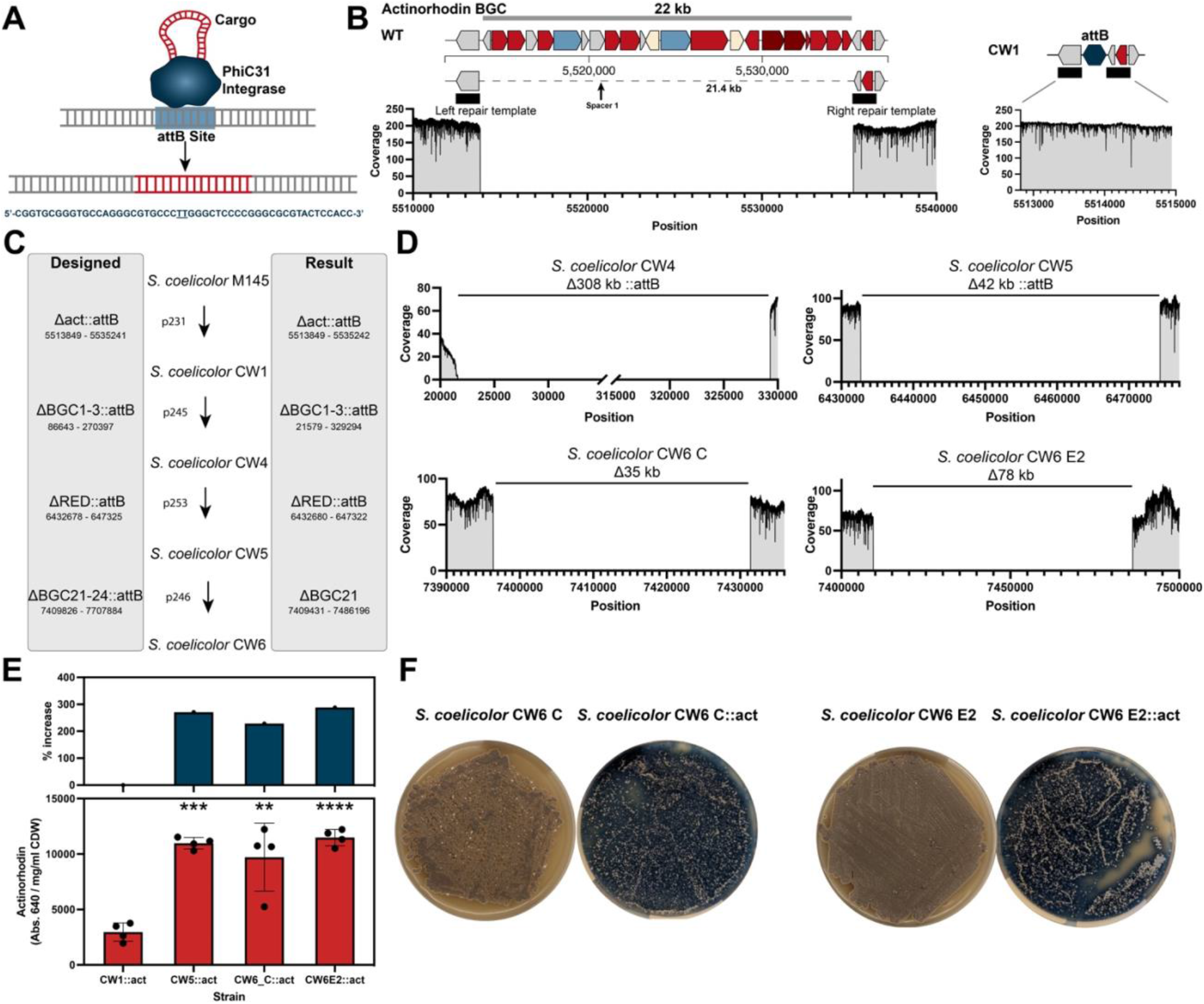
Simultaneous deletions and integrations enable streamlined genome engineering. **(A)** The PhiC31 *Streptomyces* integrase integrates cargo DNA into target attB sites. The consensus attB site from *S. coelicolor* is 51 bp long and features a central TT sequence where the cargo is integrated. **(B)** Substitution of the entire actinorhodin BGC with an additional attB site. The attB site was cloned between the repair templates. Coverage plots of mappings of ONT data against the wild type and the *in silico* generated substitution strain show precise genome engineering. **(C)** pCRISPR-Cas3 was used for construction of a *S. coelicolor* expression host using both targeted and semi-targeted deletions and substitutions. **(D)** Oxford Nanopore sequencing results for all deletions based on minimap2 mappings to the reference genome. **(E)** The final strains *S. coelicolor* CW5 and CW6 both displayed >200 % increase in actinorhodin production compared to the base strain *S. coelicolor* CW1 upon integration of an actinorhodin BGC BAC. **(F)** Phenotypes of *S. coelicolor* CW6 C and E2 without and with actinorhodin integrations. Shown in (**E**) are the means ± standard deviations of four biological replicates. Significance was tested using unpaired two tailed t-tests, where **P < 0.01, ***P < 0.001, ****P < 0.0001.

### Cas3-based genome engineering enables rapid host construction in *S. coelicolor*

Motivated by these results, we wanted to expand on this approach on demonstrate how pCRISPR-Cas3 can enable streamlined genome engineering of *Streptomyces* hosts. *S. coelicolor* has previously been engineered as a host for heterologous expression of BGCs, and a number of derivatives were constructed in the process ^43^. The most widely used ones, *S. coelicolor* M1152 and M1154 are both quadruplicate deletion strains (*Δact Δred Δcpk Δcda*) with varying additional mutations. To demonstrate how pCRISPR-Cas3 can be used for streamlined genome engineering experiments, we attempted to perform simultaneous genome reduction experiments and integrations of additional *attB* sites. It was previously shown that the integration site of BGCs can have great effects on product titers, and that integrations in the chromosomal arms, where the BGC density is bigger, generally result in higher titers^44^. Therefore, deletion of BGCs and simultaneous integration of *attB* sites in place is likely to result in desirable production phenotypes.

Based on the *S. coelicolor* M145 Δact::*attB* strain, from here on referred to as *S. coelicolor* CW1, several rounds of pCRISPR-Cas3 mediated genome engineering experiments were performed. Given the processive nature of the Cas3 nuclease, we utilized a combination of targeted and semi-targeted deletions. As observed in *S. albidoflavus* J1074 and *S. sp.* NBC1270, Cas3 can install extending deletions if larger or instable regions are targeted. We exploited that ability to not just delete single BGCs, but to also target larger regions with a high density of BGCs. The selected chromosomal regions contained antiSMASH regions 1-3 and 21-24, and encode several well characterized BGCs, including those for isorenieratene, hopene, and arsono-polyketide. Targeting of regions 1-3 using p245 resulted in an extending deletion similar to what was observed in *S. albidoflavus* J1074 and *S. sp.* NBC1270. Nonetheless, the deleted region was substituted by an additional *attB* site, suggesting that the repair templates must have been used during the recombination event. The resulting strain with four deleted BGCs and two additional attB sites was named *S. coelicolor* CW4 and used as the basis for the next round of genome engineering. Using p253, the undecylprodigiosin BGC was deleted and substituted by a third additional *attB* site. The deletion was very precise and varied only by a few nucleotides from the *in silico* designed deletion, again highlighting how pCRISPR-Cas3 can be used for both highly precise as well as semi-targeted random sized deletions. The new strain was named *S. coelicolor* CW5 and used for the final round of genome engineering, targeting regions 21-23 with p246. The resulting deletion was smaller in size than designed and resulted in only the deletion of region 21. Additionally, no *attB* site was integrated, suggesting that this deletion was fully unspecific. The final strain, named *S. coelicolor* CW6 was reduced by 6 BGCs and carries a total of four *attB* sites, three of which were installed using pCRISPR-Cas3. In total, the genome was reduced by around 450 kb, corresponding to approximately 5 % of the genome. In addition to the pCRISPR-Cas3 induced deletions, several transposition events were detected. These were relatively stable across all sequenced strains, indicating that the majority of these came already from the parental *S. coelicolor* M145 strain.

To test the constructed *S. coelicolor* CW strains, we introduced the actinorhodin BGC encoded on an integrative plasmid via conjugation into all strains. *S. coelicolor* CW1 was used as the base strain, and *S. coelicolor* CW5 and *S. coelicolor* CW6 as final strains. For *S. coelicolor* CW6, two clones were tested. In addition to *S. coelicolor* CW6 E2, in which the deletion size in region 21 was 76.7 kb, we also tested *S. coelicolor* CW6 C, where the deletion was only 35 kb in size. After conjugation, four biological replicates were selected for each strain and cultivated in ISP2 medium supplemented with 25 μg/ml of apramycin for one week in 24 well plate as described in the methods section. The actinorhodin production was measured in the supernatant at 640 nm and normalized with the cell dry weight of the respective culture. *S. coelicolor* CW5 and CW6 strains all produced significantly more actinorhodin compared to the base strain *S. coelicolor* CW1. For *S. coelicolor* CW5, the specific product yield rose from 2956 Abs._640nm_ /mg ml^-1^ CDW to 10974 Abs._640nm_ /mg ml^-1^ CDW, an increase of 271 %. For *S. coelicolor* CW6 C, the specific production was 9713 Abs._640nm_ /mg ml^-1^ CDW, corresponding to an increase of 228 % compared to *S. coelicolor* CW1. Finally, for *S. coelicolor* CW6 E2, a specific actinorhodin production of 11480 Abs._640nm_ /mg ml^-1^ CDW was measured, an increase of 288 % over *S. coelicolor* CW1. These results highlight how the streamlined deletion of BGCs and integration of additional attB sites can be used to construct potential new expression hosts.

## Discussion

Here, we demonstrated for the first-time genome engineering of streptomycetes using a type I CRISPR system. Type I CRISPR systems are the most widespread systems in bacteria. In *Streptomyces*, type I CRISPR systems are widespread, while type II systems are very rare. We therefore hypothesized that *Streptomyces* might be more amendable to genome engineering facilitated by type I CRISPR systems. The processive nature of the hallmark nuclease Cas3, which results in long ssDNA overhangs results in recombinogenic overhangs. Given that *Streptomyces* have a high homologous recombination capability^45^, we hypothesized that type I CRISPR systems are uniquely positioned to facilitate efficient genome engineering. The system used in this study is a previously characterized minimal type I-C CRISPR system^31^.

The system, called pCRISPR-Cas3, is based on our established CRISPR platform, using a pSG5 replicon-based plasmid. The CASCADE is expressed from a single promoter as a polycistronic operon. Internal RBSs facilitate successful expression of all *cas* genes. The total plasmid size of almost 13 kb is the same as those of frequently used CRISPR plasmids in *Streptomyces*, suggesting that conjugative transfer and replication in *Streptomyces* should present no problem. To overcome cloning limitations resulting from the need to clone protospacer sequences between two CRISPR repeats, as well as the absence of established type IIS restriction enzymes for high GC contexts, we designed a PCR and Gibson Assembly based cloning workflow. This workflow robustly achieved high efficiencies, however limits library-based applications due to the difficulties of cloning such with PCR based approaches. This represents an obvious improvement for future iterations of the plasmid system.

pCRISPR-Cas3 introduced highly efficient deletions of the actinorhodin BGC in *S. coelicolor*. The strong red phenotype of deletion mutants simplified screening and assessment of efficiencies. The obtained results were verified by PCR and Illumina sequencing, showing that CASCADE-Cas3 is indeed more efficient than Cas9. Previous reports of pCRISPR-Cas9 based engineering reported higher efficiencies, however were based on targeted screening of specific phenotypes ^20^. Here, we followed an untargeted screening approach. Interestingly, successful deletions with pCRISPR-Cas3 appeared to be less dependent on the selected protospacer compared to pCRISPR-Cas9. This suggests that the processive nature of the Cas3 nuclease might help to force the desired recombinations, and that Cas3 results in lower off target induced toxicity.

The longer protospacer sequence, as well as the AT rich PAM sequence, likely minimizes off-target effects, thus reducing off-target associated toxicity of pCRISPR-Cas3. Direct toxicity of pCRISPR-Cas3 was only observed when protospacers were used alone and without repair templates, forcing repair through non-homologous end joining. Unspecific deletions were only obtained when targeting the highly plastic chromosomal arms and when targeting large extending chromosomal stretches with many hypothetical and potentially essential genes. However, the large extending deletions were observed for both pCRISPR-Cas9 and pCRISPR-Cas3 and were therefore likely just the result of targeting the highly instable chromosomal arms.

pCRISPR-Cas3 was subsequently demonstrated to introduce genomic deletions in multiple model and non-model species. This suggests that pCRISPR-Cas3 can be easily implemented for genome engineering in many *Streptomyces* species. Finally, simultaneous deletions and integrations were demonstrated by integration the PhiC31 *attB* site in place of BGCs. Through multiple rounds of genome engineering, the hosts *S. coelicolor* CW5 and CW6 were constructed, which produced significantly more actinorhodin compared to the base strain *S. coelicolor* CW1 after BGC integration. The observed increase in production was likely the result of both deletion of competing BGCs, a reduced genome size, as well as multicopy integrations of the BGC. This approach is likely to enable streamlined construction of overproduction strains in several *Streptomyces* species.

Given that pCRISPR-Cas3 is based on the established pCRISPR-Cas9 plasmid system, we hypothesize that most strains in which pCRISPR-Cas9 or CRISPR-BEST were established will also be amendable to genome engineering using pCRISPR-Cas3. Likely, pCRISPR-Cas3 will become a successor to pCRISPR-Cas9 for many applications, and enable highly efficient genome engineering, including targeted deletions, random sized deletions, as well as substitutions, even in previously difficult to engineer species. By enabling highly efficient genome engineering in more *Streptomyces* species, pCRISPR-Cas3 will likely become a crucial tool for studies linking BGCs to compounds, host construction, and genome reduction studies.

## Materials and Methods

### Strains and Culture conditions

All *E. coli* work for cloning and maintenance was performed in chemically competent One Shot™ Mach1™ T1 Phage-Resistant *E. coli* cells (ThermoFisher Scientific Inc., U.S.A) cells. Strains were cultivated on LB agar plates (10 g/l tryptone, 5 g/l yeast extract, 5 g/l sodium chloride, 15 g/l agar, to 1 l with MiliQ water) or in liquid 2xYT medium (16 g/l tryptone, 10 g/l yeast extract, 5 g/l sodium chloride, to 1 l with MiliQ water) and incubated at 37 °C. If required, medium was supplemented with the appropriate antibiotics (50 ng/μl apramycin, 50 ng/μl kanamycin, 25 ng/μl chloramphenicol). *Streptomyces* strains were cultivated at 30 °C on mannitol soy flour (MS) plates (20 g/l fat reduced soy flour, 20 g/l mannitol, 20 g/l agar, 10 mM MgCl_2_, to 1 l with tap water) for spore generation. ISP2 plates (4 g/l yeast extract, 10 g/l malt extract, 4 g/l dextrose, 20 g/l agar, 333 ml tap water, 667 ml MiliQ water) were used for clean streaking. For liquid culture, 350 ml baffled shake flasks containing 50 ml of ISP2 (supplemented with the appropriate antibiotics if needed) were inoculated from spores. All shake flask cultivations were performed in shaking incubators at 30 °C and 180 rpm (311DS, Labnet International Inc., U.S.A). For actinorhodin production experiments, 24 well plates were used. 6 glass beads were added into each well, and cultivations were performed in 3.6 ml of ISP2 medium. Incubations were performed at 250 rpm and 30 °C. *Streptomyces* medium was supplemented with 12.5 ng/μl nalidixic acid and 50 ng/μl (solid medium) or 25 ng/μl (liquid medium) if required. For plasmid curing, plasmid harboring strains were cultivated at 40 °C in non-selective ISP2 liquid medium, streak on non-selective MS plates, and picked separately on selective and non-selective ISP2 plates for identification of plasmid-free clones.

### Cloning Work

All *in silico* cloning was performed in SnapGene v6.2.1 (Dotmatics Limited, U.S.A). Primers were ordered from IDT (Integrated DNA Technologies, U.S.A). The CASCADE-Cas3 operon was synthesized by GenScript Biotech Corporation and delivered as a plasmid. PCRs for cloning and of high GC *Streptomyces* elements were performed using Q5^®^ High-Fidelity DNA Polymerase with GC Enhancer (New England BioLabs Inc., U.S.A). Colony PCRs for crRNA integrations into pCRISPR-Cas3 were performed using One*Taq^®^* 2X Master Mix with Standard Buffer Enhancer (New England BioLabs Inc., U.S.A).

Minipreps were performed using NucleoSpin^®^ Plasmid EasyPure Kits (Macherey-Nagel, Germany). 1 % agarose gels were run at 100 V for 20-30 min using 6x DNA Gel Loading Dye and GeneRuler 1 kb DNA Ladder (Thermo Fisher Scientific Inc., U.S.A). Both gel purifications and PCR clean-ups were performed using the NucleoSpin^®^ PCR and Gel Clean Up Kit (Macherey-Nagel, Germany). DNA concentrations and purities were measured on a NanoDrop^TM^ 2000 (Thermo Fisher Scientific Inc., U.S.A). Restriction digestions were performed using Thermo Scientific^TM^ FastDigest enzymes. Differing from the manufacturer’s recommendations, the restriction digests were performed at 37 °C with extended incubation times of 2 hours, followed by inactivation for 10 min at 75 °C. 5 U/µL T4 DNA Ligase (Thermo Fisher Scientific Inc., U.S.A) was used for ligations overnight at room temperature. Gibson Assemblies were performed using NEBuilder^®^ HiFi DNA Assembly Master Mix (New England BioLabs Inc., U.S.A) following the manufacturer’s instructions. Sequence verifications were performed by Sanger sequencing using Mix2Seq overnight kits (Eurofins Genomics Germany GmbH, Germany).

### Genome Mining & Protospacer Prediction

Prediction of biosynthetic gene clusters was performed using antiSMASH 6.0.1 ^46^. Using the antiSMASH output, protospacers were predicted for pCRISPR-Cas9 based deletions using CRISPy-web ^47^. Protospacers for pCRISPR-Cas3 were identified by searching for 5’-TTC-N_1_-N_8_-3’ sequences within the target region with no mismatches in the genome. The “find similar DNA sequences” function of SnapGene was used to access potential off-targets with mismatches. The endogenous CRISPR system in *S. albidoflavus* J1074 was identified by searching CRISPRCasdb ^48^. The presence of predicted CRISPR arrays and Cas genes was subsequently manually verified in the sequenced strain of *S. albidoflavus* J1074.

### Interspecies Conjugations

Transfer of plasmid DNA into *Streptomyces* strains was performed through interspecies conjugations. Spores for conjugations were prepared as described in Tong et al. ^35^.

Homemade chemically competent or room temperature competent^49^ *E. coli* ET12567 pUZ8002 ^50^ cells were transformed and plated on LB plates supplemented with 50 ng/μl apramycin, 25 ng/μl chloramphenicol, and 50 ng/μl kanamycin. All transformants were washed off using LB medium and 5 ml LB overnight cultures inoculated with 50 μl. 2 ml of the overnight cultures were harvested the next day and washed twice at 2000xg with 1 ml of fresh LB medium. 500 μl of resuspended *E. coli* ET12567 pUZ8002 cells were then mixed with 500 μl of filtered spore suspension and spread on MS + 10 mM MgCl_2_ plates. The next day, overlays were performed with 1 ml of ddH_2_O and 5 μl of 50 mg/ml apramycin. Exconjugants were picked using wooden toothpicks and transferred to selective ISP2 plates supplemented with nalidixic acid.

### Streptomyces Colony PCRs

For colony PCRs on *Streptomyces* colonies, pieces of the colonies were scraped off the plates and transferred to PCR tubes containing 50 μl of 10 % DMSO. The tubes were sealed and boiled for 10 min at 99 °C, transferred to dry ice for 20 min, and then again boiled for 10 min. The entire process was repeated one more time, and the tubes then spun down to separate the debris from the supernatant. 1 μl of the supernatant was used as template for colony PCRs. All *Streptomyces* colony PCRs were performed using Q5^®^ High-Fidelity DNA Polymerase with GC Enhancer (New England BioLabs Inc., U.S.A).

### Whole Genome Sequencing & Bioinformatic Analysis

Genomic DNA was isolated from *Streptomyces* plates or liquid cultures using the DNeasy PowerLyzer PowerSoil Kit (Qiagen, Germany). The bead beating was performed using a TissueLyser LT (Qiagen, Germany) at default settings for 7 mins. Quality control was performed using a NanoDrop^TM^ 2000 for purity and Qubit 4 Fluorometer for concentration measurements (Thermo Fisher Scientific Inc., U.S.A). Genomic DNA was further run on 0.8 % agarose gels to assess fragmentation. Illumina sequencing was performed by Novogene Co., Ltd (Bejing, China). Libraries were prepared using the NEB Next® UltraTM DNA Library Prep Kit (New England Biolabs, U.S.A) with a target insert size of 350 nt and six PCR cycles. trimgalore (https://github.com/FelixKrueger/TrimGalore) and breseq 0.33.2 ^51^ were used for read trimming and data analysis of illumina sequencing data. The following commands were used:

*trim_galore -j 8 --length 100 -o illumina --paired --quality 20 --fastqc --gzip file(s)*
*breseq -r (full genetic background reference).gb trimmed_1.fq.gz trimmed2.fq.gz -j 12*

Using samtools^52^, the read coverage was extracted from the the breseq output as follows:

*samtools depth -a sample.bam > sample.coverage*

Oxford Nanopore sequencing was performed as described by Alvarez-Arevalo et al. ^53^ and basecalled using Guppy (5.0.17+99baa5b, client-server API version 7.0.0 or 6.3.8+d9e0f64, client-server API 13.0.0) applying the high-accuracy model and excluding reads shorter than 1 kb.

Minimap2 2.18 ^54^was used to map reads and samtools was used to extract the reads as follows:

*minimap2 -a reference.fa reads.fastq.gz > mapping.sam*
*samtools sort mapping.sam > mapping.sam.sort.bam*
*samtools depth -a mapping.sam.sort.bam > sample.coverage*

Mappings were visualized using Artemis ^55^. For plotting, the regions of interested were extracted from the .coverage files and plotted in Prim 9 (GraphPad Software, U.S.A). *De novo* assemblies were performed using flye 2.9 ^56^. Assembly graphs were visualized using bandage 0.8.1^57^.

### Cell Dry Weight Measurements

The endpoint cell dry weight was measured during harvesting of cultures. 2 ml tubes were dried overnight at 60 °C. For each strain, three replicates were measured. The tubes were weighed and numbered. 2 ml of cell culture were added, spun down at max. speed, and washed with ddH_2_O. After removing all supernatant, the tubes were dried overnight at 60 °C and measured the next day on a Sartorius Qunitix scale (1 mg). The weight of the specific empty tube was subsequently subtracted to give the cell dry weight.

### Actinorhodin Measurements

For actinorhodin quantification, 2 ml of shake flask cultures were harvested by centrifugation for 10 min at max. speed. The supernatant was transferred to new tubes, and 100 μl were transferred to F-bottom, clear, 96 well microplates (Greiner Bio-One International GmbH, Austria). 50 μl of 3 M NaOH were added to each well and carefully mixed by pipetting up and down. The samples were then measured in a BioTek EPOCH2 microplate reader (Agilent Technologies, Inc.) at 640 nm.

## Data Availability

All whole genome sequencing data is available in the sequence read archive under BioProject PRJNA966932. Additional supplementary information such as plasmid and primer lists are available at: https://figshare.com/s/e1bfb388df0ae502034e or doi: 10.11583/DTU.22786157

## Acknowledgments

This work was supported by grants of the Novo Nordisk Foundation (NNF16OC0021746, NNF20CC0035580).

## Author Contributions

C.M.W. and T.W conceived and design the project. C.M.W, P.G, T.G, and R.S. carried out laboratory experiments. C.M.W analyzed data. D.F. analyzed Oxford Nanopore sequencing data. C.M.W wrote the manuscript with input from all authors.

