## Supplementary Information for "CASCADE-Cas3 Enables Highly Efficient Genome Engineering in *Streptomyces* Species"

Supplementary Information Figure 1

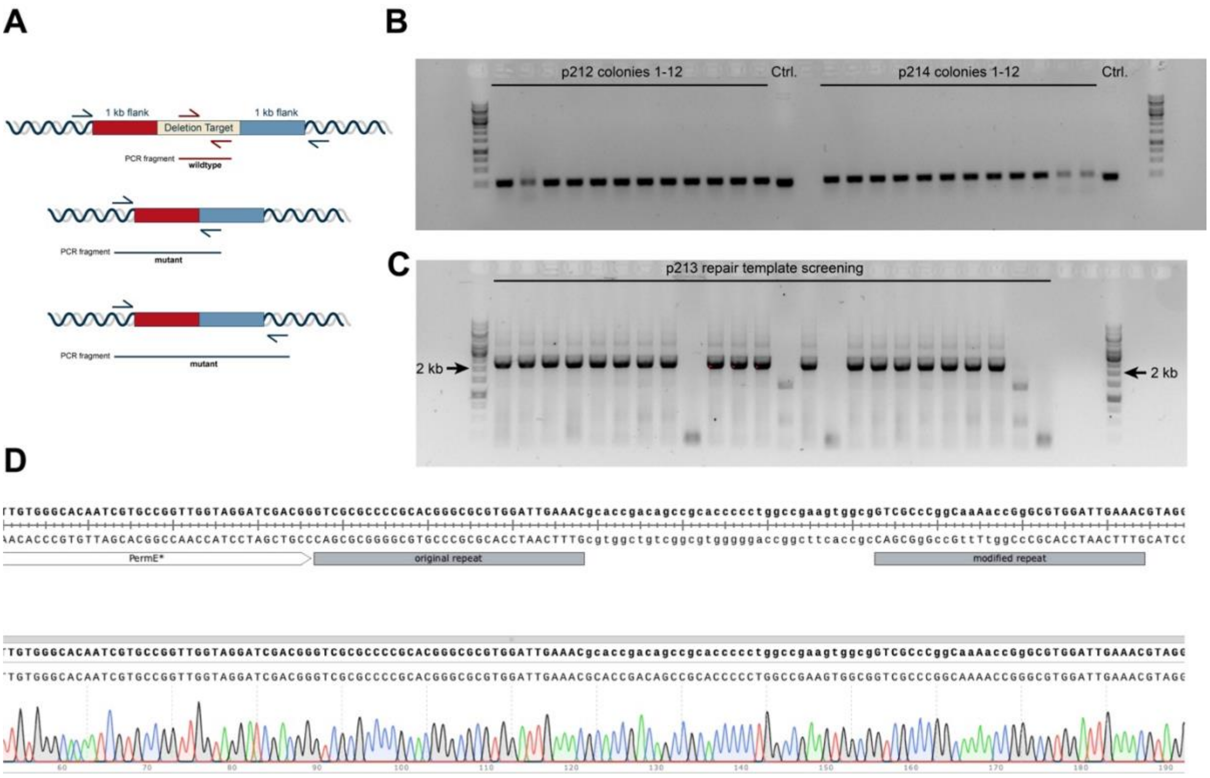


***Supplementary Figure 1:*** **A)** Schematics for different screening approaches. Colony PCRs can be performed using primers binding inside the deletion region for screening of drop out mutants, pairs of primers with one binding inside the repair templates bridging the junction site and one primer binding in the chromosome outside, or with two primers both outside of the repair templates. Given the challenges associated with obtaining large high GC PCR fragments, the second approach is most widely used. **B)** Exemplary results for colony PCRs for protospacer (top) and repair template (bottom) integrations. The differences between the empty pCRISPR-Cas3 repeat region and the same region with an integrated protospacer is large enough to differentiate negative colonies from positives, as these lead to slightly higher bands in the gel. **C)** Example of an alignment of Sanger sequencing data of a cloned protospacer.

### Supplementary Information Figure 2

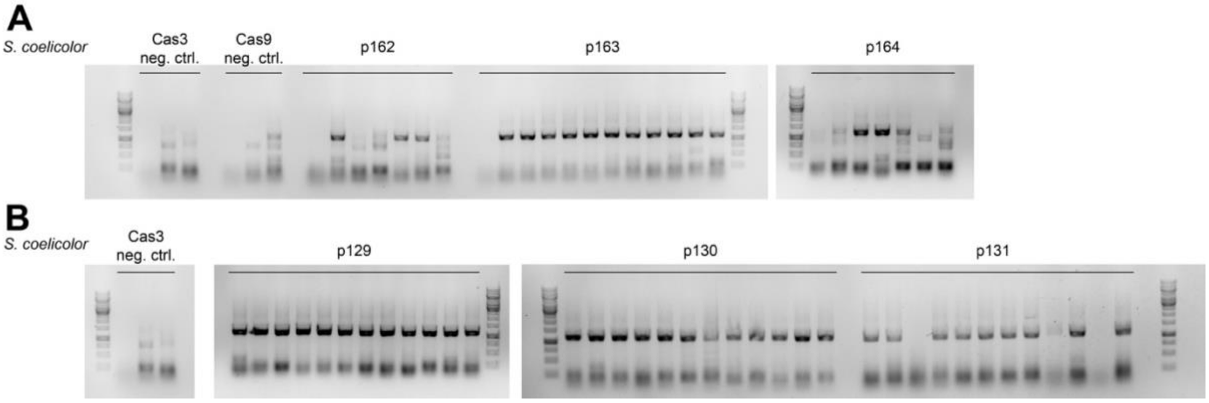


***Supplementary Figure 2:*** **A)** Gel pictures of colony PCR screening for pCRISPR-Cas9 mediated deletions of the actinorhodin biosynthetic gene cluster in *S. coelicolor* M145. p162-p164 carry the “left”, “middle”, and “right” protospacer, respectively. One of the negative controls produced a light bands also around 1 kb in size, however upon closer inspection that band sits slightly higher than the expected band for deletions. Positive samples produced strong bands slightly lower than the unspecific light band. **B)** Colony PCRs for pCRISPR-Cas3 mediated deletions of the actinorhodin biosynthetic gene cluster. Strong bands slightly above the 1 kb ladder were observed for almost all colonies. p129-p131 carry the “left”, “middle”, and “right” protospacers, respectively.

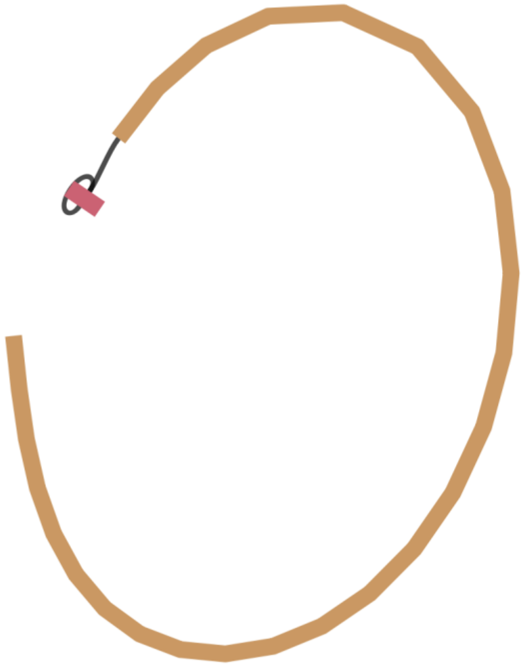

***Supplementary Figure 3:*** Assembly graph of a *de novo* assembly of NBC1270 after targeting BGCs 1-3 with pCRISPR-Cas9. The assembly was visualized using bandage. The graph highlights how a sequence stretch at the far chromosomal end was replicated multiple times. Based on the coverage, around 12 replications are expected.

**Supplementary Information Table 1:**

Blast results for *Salb*Cas3 and *Spy*Cas9. Can be downloaded here: https://figshare.com/s/e1bfb388df0ae502034e or doi: 10.11583/DTU.22786157

**Supplementary Information Table 2:**

Cblaster output for *Salb*CASCADE. Can be downloaded here: https://figshare.com/s/e1bfb388df0ae502034e or doi: 10.11583/DTU.22786157
